# On the correlation between material-induced cell shape and phenotypical response of human mesenchymal stem cells

**DOI:** 10.1101/2020.05.13.093641

**Authors:** Aliaksei S Vasilevich, Steven Vermeulen, Marloes Kamphuis, Nadia Roumans, Said Eroumé, Dennie G.A.J. Hebels, Jeroen van de Peppel, Rika Reihs, Nick R.M. Beijer, Aurélie Carlier, Anne E. Carpenter, Shantanu Singh, Jan de Boer

## Abstract

Learning rules by which cell shape impacts cell function would enable control of cell physiology and fate in medical applications, particularly, on the interface of cells and material of the implants. We defined the phenotypic response of human bone marrow-derived mesenchymal stem cells (hMSCs) to 2176 randomly generated surface topographies by probing basic functions such as migration, proliferation, protein synthesis, apoptosis, and differentiation using quantitative image analysis. Clustering the surfaces into 28 archetypical cell shapes, we found a very strict correlation between cell shape and physiological response and selected seven cell shapes to describe the molecular mechanism leading to phenotypic diversity. Transcriptomics analysis revealed a tight link between cell shape, molecular signatures, and phenotype. For instance, proliferation is strongly reduced in cells with limited spreading, resulting in down-regulation of genes involved in the G2/M cycle and subsequent quiescence, whereas cells with large filopodia are related to activation of early response genes and inhibition of the osteogenic process. Thus, we have started to unravel the open question of how cell function follows cell shape. This will allow designing implants that can actively regulate cellular, molecular signalling through cell shape. Here we are proposing an approach to tackle this question.

## Introduction

Cells adapt to new situations by perceiving information, and transforming it into changes in their gene and protein expression repertoire (Tosh & Slack, 2002). Typical examples are diffusible morphogens (Potter, 2007) that control stem cell differentiation during development and regeneration (Hu et al., 2015; Liddiard & Taylor, 2015), UV exposure resulting in the synthesis of pigment skin melanocytes (Lin & Fisher, 2007), lack of oxygen initiating the HIF1 mediated hypoxia response (Semenza, 2001) and shear forces in blood vessels regulating endothelial cell physiology (Hahn & Schwartz, 2009). Adaptation to the environment can also materialize as change in cell shape (Nelson, 2003). Elongated multinuclear muscle cells are particularly well suited to exert stretching forces (Cui et al., 2015), whereas the globular shape of lipid cells is optimized for volume-efficient fat storage (Chen et al., 2016).

Cell shape thus follows cell function, but is cell shape also important to control cell function? Does cell function follow cell shape? Loss of physiological cell shape is sometimes accompanied by pathology (Cook, Hardy, McConnaughey, & Zuker, 2008). For instance, in tendinopathies loss of extracellular matrix integrity in tendon and ligaments is preceded by a change from the typically elongated tenocyte to a more rounded shape (Gehwolf et al., 2016). Similarly, cancer cell malignancy is often preceded by a change in cell morphology (Lee, Abdeen, Wycislo, Fan, & Kilian, 2016), suggesting that shape changes may be the cause rather than the consequence of the functional change. Experimental evidence for “function follows shape” comes from in vitro experiments where cell shape can be controlled using micropatterning techniques. Basic cellular decisions such as differentiation, apoptosis or metabolic rate can be controlled by merely controlling cell shape (McBeath, Pirone, Nelson, Bhadriraju, & Chen, 2004), suggesting that cell shapes generate signals which are transduced into the nucleus and result in changes in gene and protein expression (Vogel & Sheetz, 2006).

An important source of cell shape signaling is the actin cytoskeleton which changes upon shape change, leading to changes in transcriptional activity of transcription factors such as MRTF, SRF, YAP and EGR (Chiquet, Gelman, Lutz, & Maier, 2009; Papachroni, Karatzas, Papavassiliou, Basdra, & Papavassiliou, 2009; Sun, Guo, & Fässler, 2016). However, nature has more mechanosensitive proteins in store. Stretch-activated ion channels open upon stretching of the plasma membrane resulting in the influx of ions which trigger several signalling cascades (Coste et al., 2010). Many proteins contain mechanosensitive protein domains, e.g. the focal adhesion protein talin (Roca-Cusachs, Gauthier, Del Rio, & Sheetz, 2009), and some proteins contain a so-called BAR domain which is able to sense the curvature of a membrane and transduce that into a signal (Wiche, Osmanagic-Myers, & Castanon, 2015). The generation of cell signalling events under the control of shape can be clustered under the term mechanotransduction (Iskratsch, Wolfenson, & Sheetz, 2014), and an open question in this field is how many different mechanobiological signals can be picked up by how many molecular circuits. The analogy is that of the canonical signalling pathways, where a dozen well-described families of diffusible molecules (hormones, metabolites, etc.) bind to and activate their cognate families of receptors and initiate signalling cascades.

We aim to identify how many mechanobiological pathways exist and how they are activated by cell shape parameters. How specific are the mechanobiological pathways and can we correlate shape features to gene expression and changes in phenotype? To answer these questions, we (Unadkat et al., 2011) and others (Masaki, Schneider, Zaharias, Seabold, & Stanford, 2005) have used surface topography to control cell shape and direct cell function, because shape mediated control can be applied to functionalize the surface of medical devices. In this study, we have generated a library of topography-induced cell shapes and were able to couple shape induced cell signalling to several different phenotypes, including molecular and morphological (Figure 1a). We used bone marrow-derived mesenchymal stem cells, because they are multipotent, their progeny has very distinct differences in cell shape and there is ample evidence for shape-directed differentiation.

**Figure 1.**
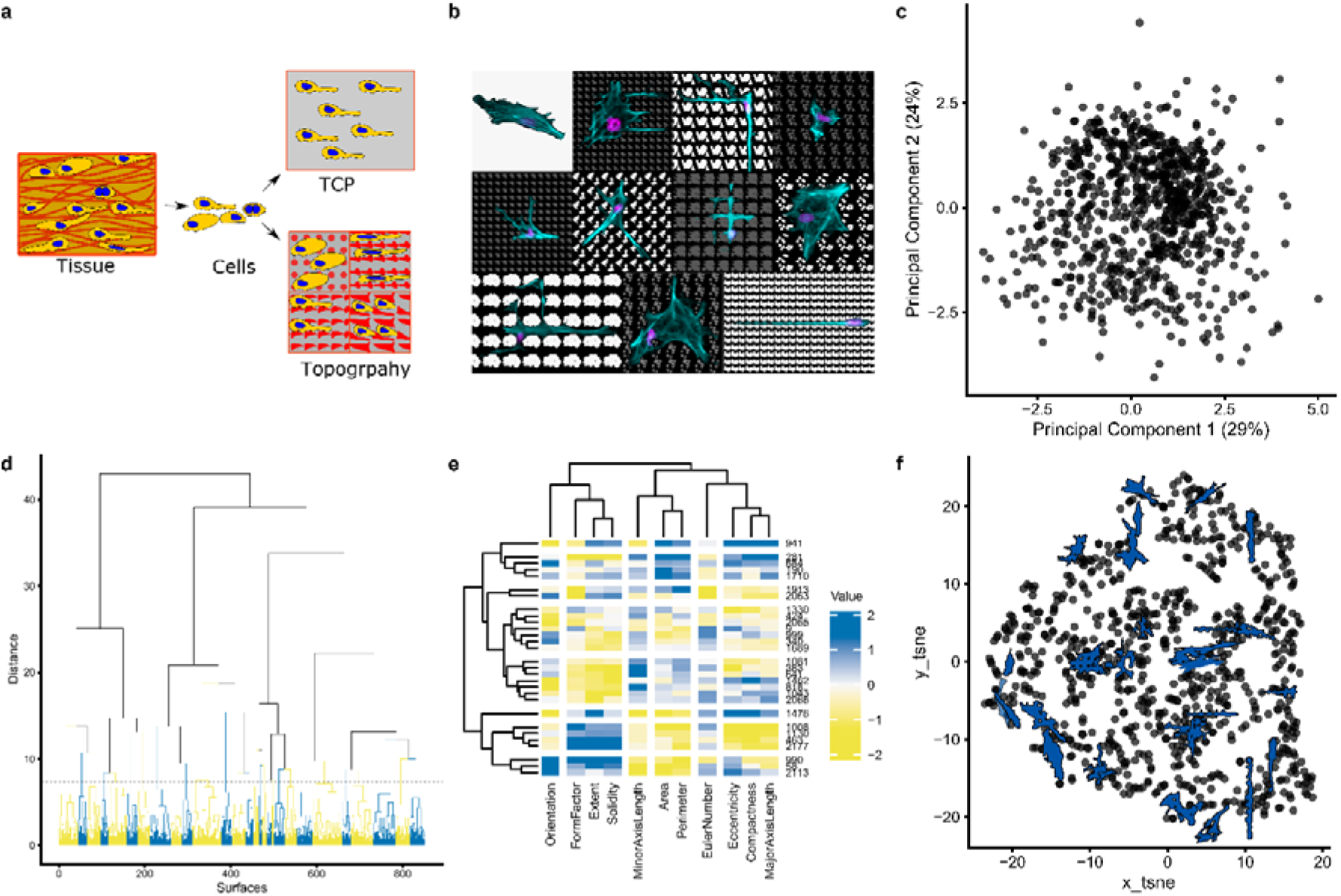
Clustering of cell shape diversity. **A.** Surface topography as a model system to investigate cell shape-induced cells response. TCP: tissue culture polystyrene. **B.** The montage image that represents the diverse cell of hMSCs seeded on different topographies and flat surface (upper, left). Nuclei were stained with DAPI (magenta), actin was stained with phalloidin (cyan). **C**. Principal components analysis of all cell shapes on the TopoChip. The first two principal components are shown. Note that no noticeable clusters are observed. **D.** Cluster gram representing a clustering of TopoChip cell shape data. Blue and yellow colours distinguish different clusters. **E.** Heatmap representing the median values of cell shape features on the chosen medoid topographies **F.** tSNE (t-distributed stochastic neighbor embedding) visualization of the obtained clusters. Cells shape silhouettes on 28 selected surfaces are represented.

## Results

### Cell shape diversity can be captured by feature-based clustering of a library of cell shapes

To systematically explore the relation between cell shape and phenotypic response, we exposed human bone marrow-derived mesenchymal stem cells (MSC) to a library of 2176 unique topographies (TopoChip) and observed a plethora of different cell shapes (Hulshof et al., 2017) which were distinct from the MSCs cultured on the flat substrate (Figure 1b). For example, we observed cells with an elongated narrow cell body without any protrusions which resemble the shape of tenocytes (Figure 1b, bottom-right cell) or had the cuboidal shape typical for bone lining cells; still, others appeared neuronal. Thus, topographies are able to induce cell shapes, some of which reminisce different mesenchymal cells *in vivo*.

To capture shape diversity, we generated a morphological fingerprint for each cell on each topography using 11 shape descriptors including Area, Perimeter, Eccentricity and Form Factor (Carpenter et al., 2006) (Supplementary Tables 1, 2). We pruned the cell database to remove outliers and imaging artefacts and focused on topographies that consistently induce the same cell shape (Supplementary Materials and Methods). The resulting selection of 851 cell shapes was plotted by Principal Components Analysis (PCA, Figure 1c), which reduces data dimensionality and allows identifying cells with similar shapes. We observed a continuum of cell shapes with some densely and sparsely occupied areas, demonstrating that some shape features are more abundant in our shape collection than others. This was confirmed by hierarchical clustering, where the tree structure displays four distinct branches, which are further divided until it reaches single cell shapes (Figure 1d). For further phenotypic experiments, we sampled the medoid surfaces from 28 clusters, which together span the whole range of shape feature range, and reproduced them on 12 mm polystyrene disks. Visual inspection of the MSCs again shows a large diversity of cell shape, which is further highlighted by their cell shape fingerprints (Figure 1e). The 28 surfaces are thus representing cell shape diversity induced by the TopoChip library (Figure 1f) and further permits well plate compatible biological assays.

### Shape features of mesenchymal stem cells correlate to their phenotypic response

Cell shape is related to cell function (Etienne-Manneville & Hall, 2002; Fletcher & Mullins, 2010; Ron et al., 2017), so we probed the relation between cell shape features and a series of basic cell phenotypes, i.e. proliferation, apoptosis, protein biosynthesis, migration and differentiation, in MSCs exposed to the 28 groups of topographically-patterned surfaces and to flat control surfaces. We found that topographies induced profound differences in all phenotypic assays, demonstrating that basic cell biological processes are influenced by surface topographical cues. For instance, we observed a big difference in the rate of proliferation, ranging from 12% to 59% of EdU-positive cells, between the low and high scoring surfaces (Figure 2a). Differentiation was also affected, with, e.g. a six-fold difference in lipid production under adipogenic conditions between the lowest and the highest performing surface (Figure 2b). None of the surfaces exclusively affected one biological property, yet a comparison of the magnitude of the phenotypic response showed that each surface induces a distint phenotypic fingerprint (Figure 2c, Supplementary Figure 1). This is further illustrated in Figure 2d, in which we clustered the surfaces based on phenotypic response. Flat surfaces formed a distinct cluster, demonstrating that cells on topographies share part of their phenotypic response which is different from the behaviour of cells on a flat surface (Figure 2d). A cluster was formed by surfaces 2063 and 1130, located between the flat surface and the other topographies. Interestingly, the topographies on these surfaces are composed of sparsely located pillars, which resemble the flat surface, inducing relatively similar cell shape and similar phenotypic scores.

**Figure 2.**
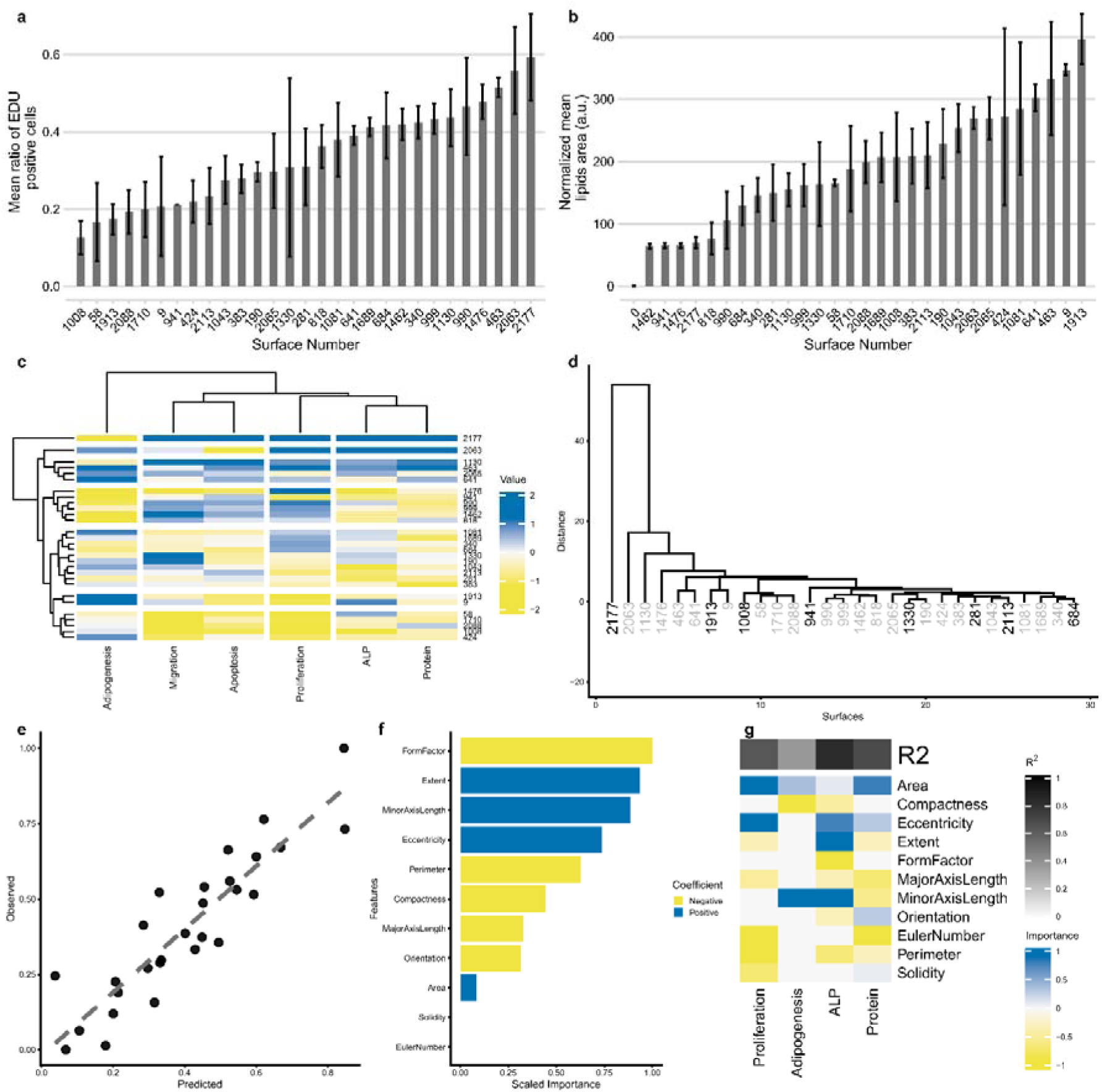
Profiling of phenotypic responses of hMSCs to topographies. **A.** Ranking of surfaces based on the percentage of EdU staining. hMSCs were starved for 24 hours, seeded on topographies overnight, and EdU was added 48 hours prior to fixation. The bar represents mean of up to 3 replicas; error bar represents standard deviation. **B.** Ranking of surfaces based on area covered by lipids. hMSCs were cultured on topographies in adipogenic medium for three weeks, and lipids were stained with Oil red O. Bar represents mean of up to 3 replicas, error bar represents standard deviation. Surface number 0 represents hMSCs on a flat surface in control media. **C.** Heatmap represents the performance of the 28 selected topographies in 6 functional assays. **D.** Clustering of the surfaces based on the readout in the phenotypic assays. Surfaces highlighted with black colour were selected for microarray analysis. On top, goodness of Fit R2 values reported. **E.** Performance of the Lasso regression model on predicting ALP induction based on the 11 cell shape features. **F.** Cell shape features that are important for predicting ALP induction. **G.** Importance of the features for predicting the outcome of the phenotypic assays obtained with Lasso model.

We assume that the phenotypic fingerprint is the result of the specific combination of multiple cellular signals induced by the topographies, some of which are directly related to cell shape, but others may not. For instance, it is conceivable that the direct effect of topography on nuclear shape influences gene expression and thus phenotype (Anselme, Wakhloo, Rougerie, & Pieuchot, 2018), whereas we did not include nuclear shape features in our analysis. To assess the hypothesis that phenotypic responses are directly related to cell shape, we constructed computational models using the Lasso algorithm in which cell shape features were correlated to each specific phenotype.

The most accurate computational model was able to predict alkaline phosphatase (ALP) expression, a marker for osteogenic differentiation, with high accuracy (goodness of fit of 0.82, Figure 2e). The shape parameters FormFactor and Minor Axis length positively correlate with ALP expression, whereas Extent showed a negative correlation (Figure 2f). Thus, a typical ALP positive cell has an elongated, irregular shape with many protrusions, which is consistent with previous work (Hulshof et al., 2017) (Figure 3a).

**Figure 3.**
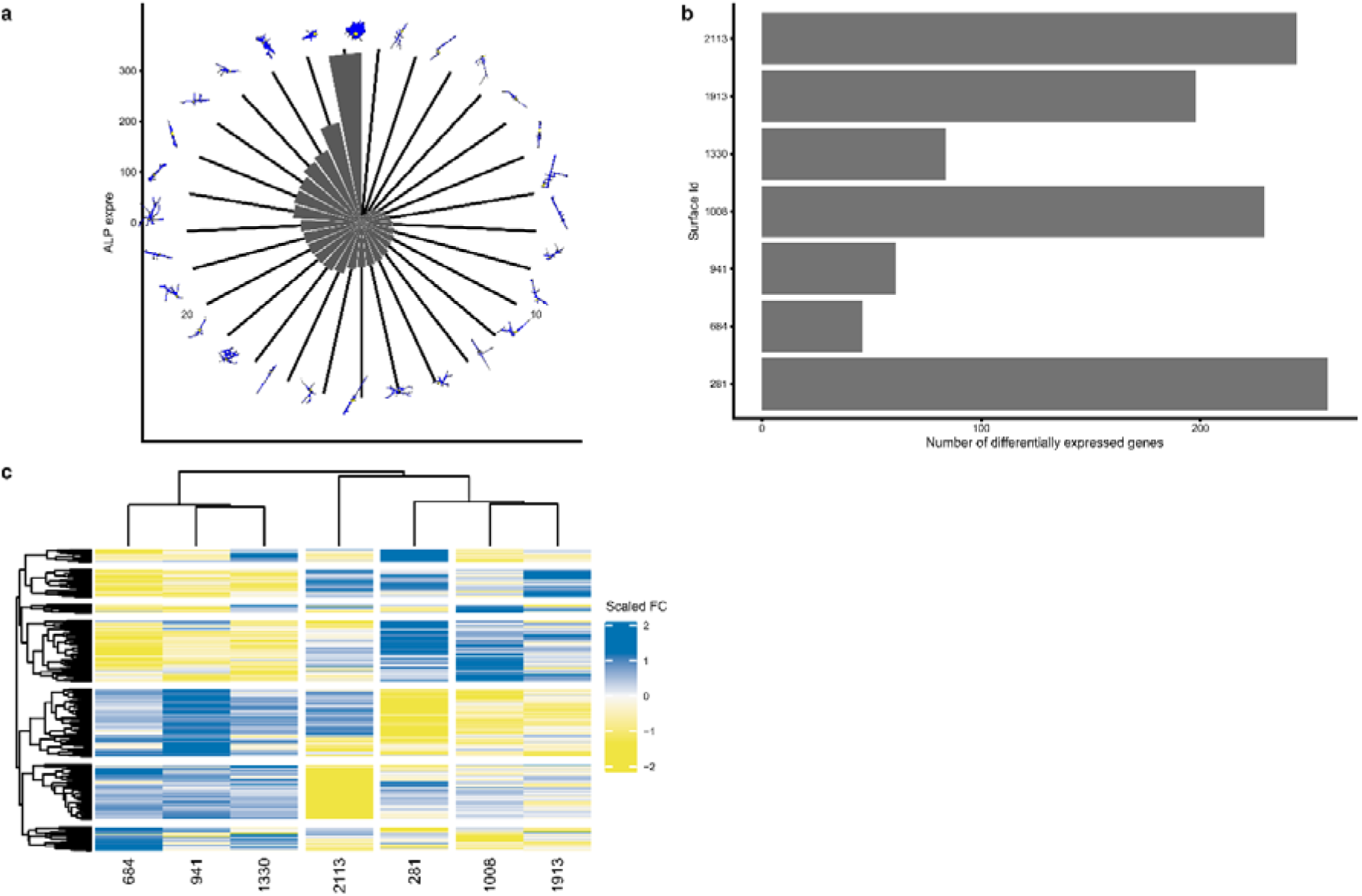
Gene expression on selected topographies. **A.** Polar Bar plot that represent level of ALP expression on different topographies, ordered from lowest to highest. On top, representative cell shapes silhouettes of the hMSCS cultured on the surface topographies are shown. **B.** A number of differentially expressed genes per topography. For identification of Differentially expressed genes (DEGs), we used data from a flat surface as a reference. **C.** Heatmap represents a unique gene expression signature of cells on each topography. To express surface-specific DEGs Fold Change (FC) values were scaled by subtracting mean and dividing by sd of DEG FC for seven surfaces. (standard scaling approach).

The second most accurate computational model predicted protein biosynthesis with goodness of fit 0.68, in which Area shows a positive correlation, whereas Euler Number (the number of the “holes” in the object, which can appear when the cell is forming a protrusion around a pillar), negatively correlates to protein synthesis (Supplementary Figure 2). We were also able to produce predictive models for cell proliferation (goodness of fit 0.63) and adipogenesis (0.37) but were unable to construct a model to predict apoptosis and migration (Figure 2g). This suggests that parameters other than explaored shape descriptors are responsible for differences in those readouts.

To summarise, surfaces selected based on the diverse cell shapes also induced diverse cell response in functional assays. Investigated cell shape descriptors could partially model the phenotypic variance. Different shape features account for different phenotypes, this suggests that shape affects the phenotype through various molecular mechanisms.

### Topographies induce distinct but overlapping gene expression profiles in MSCs

To assess the landscape of genes under the control of surface topography, we exposed MSCs to seven topographies that span the phenotypic space and to flat control surfaces and assessed the transcriptome after 24 hours. We observed between 46 to 258 differentially expressed genes with a fold-change > 1.5 as compared to flat (Figure 3a). Each topography induced a unique fingerprint at the gene expression level (Figure 3b), but with various degrees of overlap. Differential gene expression data was used to identify molecular pathways which were activated. Lists of Differentially Expressed Genes (DEGs) per topography were queried in the Connectivity Map (CMap) (Subramanian et al., 2017), a library of gene expression signatures induced by chemical compounds or genetic interference (perturbants), and connectivity scores between the topography-induced DEGs and Connectivity Map DEGs were retrieved. We focused on perturbant classes (PCL), which are groups of signatures in which members belong to the same gene family or are targeted by the same compound (Figure 4a). Some PCLs were shared by most topographies such as perturbants that induce *Cell cycle inhibition gain of function* or known inhibitors such as *JAK inhibitor* and *IKK inhibitors*, demonstrating that some signaling events are always induced when an MSC encounters a topography. In agreement with this, proliferation is impaired on all topographies relative to flat. Other PCLs were present in only one or a few topographical gene lists such as *LIM class homeoboxes GOF* or *glycogan synthase kinase inhibitor*, suggesting that some topographies induce distinctive signaling events.

**Figure 4.**
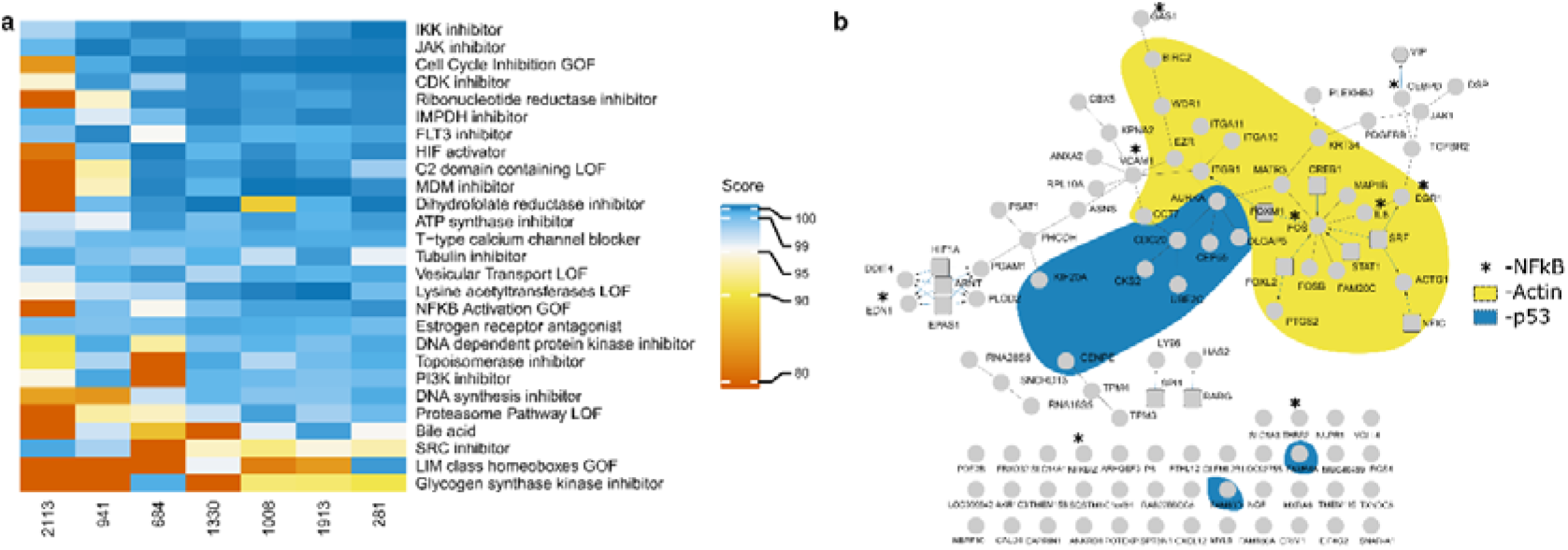
Network of genes and cell biological processes regulated on seven distinct topographical surfaces. **A.** Gene expression in hMSCs on seven different topographies was compared to hMSCs on a flat surface and differentially expressed genes were compared in the Connectivity Map, a gene expression database with more than 1 million profiles. All processes that have absolute score value above 99 at least for one condition are depicted, with 0 low and 100 as high similarity. Biological processes have been ranked based on the number of topographies that can affect the process (specificity). **B.** Gene expression in hMSCs on seven different topographies was compared to hMSCs on a flat surface, and 437 differentially expressed genes with an adjusted p-value of less than were detected. The list was uploaded in ConsensusPathDB, which retrieves correlations between genes and proteins from a number of databases. Circular nodes in the graph represent the genes; rectangles are genes retrieved from a transcription factor database, thus transcriptional control. Edges represent reported relationships. The yellow and grey shading represent two clusters of genes with published involvement in actin related-processes and p53 respectively, based on literature study. Each circular node is a pie chart indicating in which topographies the gene is differentially expressed.

To investigate how the DEGs collaborate to elicit a phenotypic response, we selected all DEGs absolute value with fold change >1.9 from all seven topographies and created a gene network (Figure 4b). Genes, represented by nodes, are connected by edges based on binary protein interactions described in public resources (such as KEGG, Reactome, and WikiPathways) using ConsensusPathDB software (Kamburov et al., 2010). 51 out of 91 genes were placed in this topography-induced gene network, suggesting an extensive overlap in functionality. Based on Gene Ontology and manual literature research, we noticed three main clusters of cellular processes, i.e. related to actin, p53 and NF-□B signaling, respectively. Further literature research revealed that many DEGs in our network are specifically expressed during the transition from the G2 to M phase of the cell cycle (e.g. Aurora Kinase and CDC20) and are repressed in cells grown on topographies. We noticed a significant overlap in topographical DEGs and those associated with the p53/DREAM complex, which regulates quiescence (Supplementary Figure 3). We previously reported on topography-induced quiescence as a typical topography-induced phenotype (Beijer et al., 2019). Whether actin, NF-kB and proliferation are in the same signalling network or represent separate signalling pathways induced by different cell shapes remains to be investigated.

### Shape-dependent phenotypic responses show a strong association with specific genes

To determine the correlation between gene expression, cell shape and phenotype, we calculated pairwise Spearman correlations between cell shape descriptors, gene expression, and phenotype score levels on all topographies. Many genes correlate to shape features such as Solidity and Perimeter (Figure 5a), whereas far fewer genes correlate to Compactness or EulerNumber. We also noted that relatively few genes correlated with protein synthesis (Figure 5b), ALP expression was positively correlated with many genes, and Adipogenesis negatively correlated with gene expression. We were particularly interested in situations where the Gene-Shape-Phenotype triad of correlations was robust and consistent (Figure 6a). For example, Spearman correlation between VCAM-1 gene expression and hMSC proliferation was -0.75 (Figure 6b), and at the same time, Spearman correlation between cell Area and proliferation was -0.71 (Figure 6d), as the result the Spearman correlation between Cells Area and VCAM-1 was 0.71 (Figure 6f). This implies that topographies that promote large cell areas are associated with a high level of proliferation and the expression of the VCAM-1 gene. Another clear example is the strong correlation between expression of the transforming growth factor receptor 2 (TGFBR2), ALP protein expression and Extent with Spearman correlation values, correspondingly, - 0.96, 0.67, -0.75 (Figure 6 c,e,g). Interestingly, a strong association between ALP protein levels and TGFBR2 are reported in the literature (MS Castro-Raucci et al., 2016). Similar to what we found here, in a previous study we discovered a strong association between the amount of protrusions and ALP protein level (Hulshof et al., 2017).

**Figure 5.**
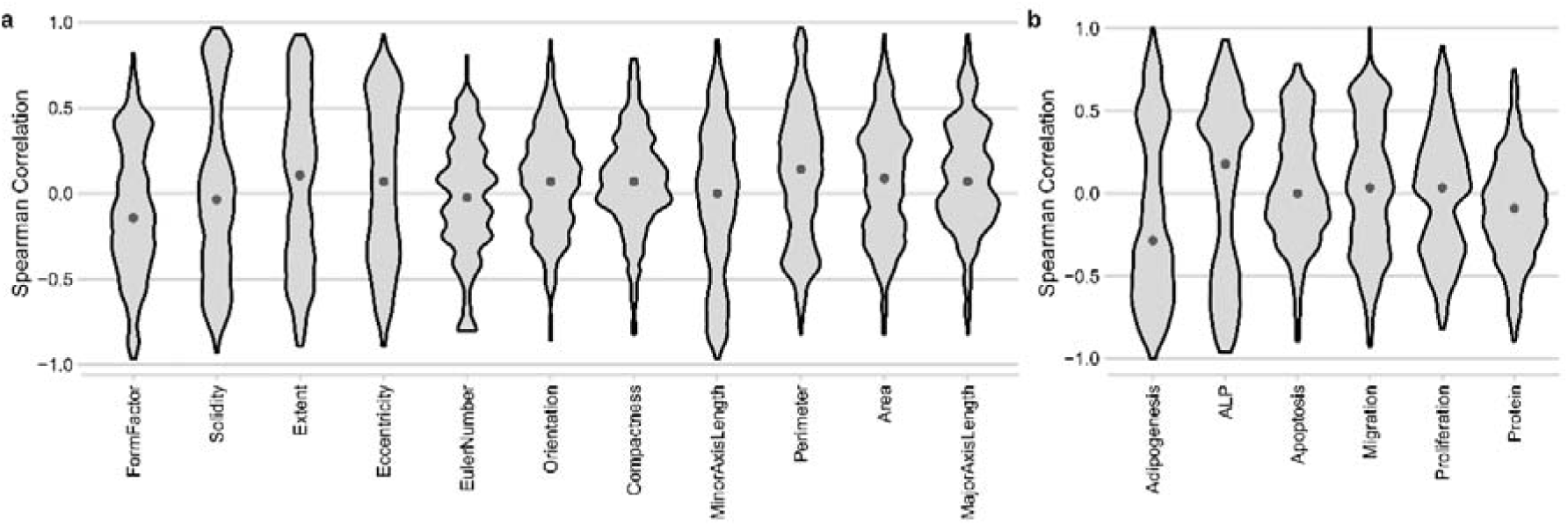
Correlation of gene expression to shape features or phenotype. Spearman calculated for all different samples a combined in 1 plot in which the distribution of outcomes can be observed. Violin plot representing the distribution of genes’ Spearman correlation to either a shape feature (**A**) or a phenotype (**B**). Each gene was plotted against the respective feature and correlation presented as a Spearman score. Solidity has many high and low scoring correlation values, meaning that Solidity correlates strongly with gene expression. Most Spearman score for Euler number is close to zero, meaning that few genes’ expression strongly correlate to this feature.

**Figure 6.**
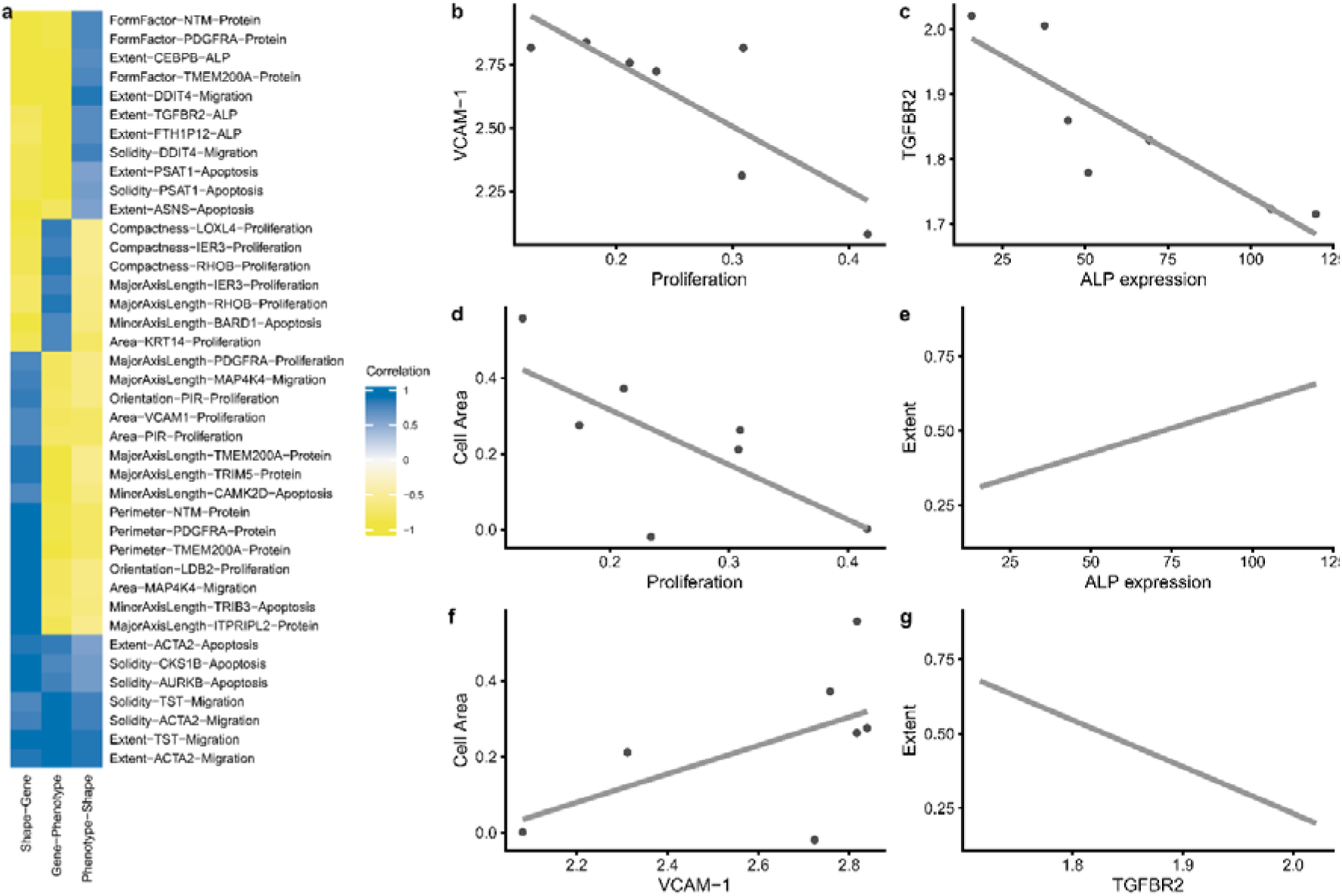
Correlation between shape feature-gene expression and outcome in the phenotypic assay. **A.** Gene expression, cell shape and phenotypic data from hMSCs grown on the seven selected surfaces were compared by Spearman correlation between the 11 shape features, 437 genes and six phenotypes. Top three correlations per unique combination Shape parameter-phenotypic assay with the highest positive or negative values are shown. **b**,**c**,**d**,**e**,**f g** shows corresponding scatter plots with raw data.

Browsing the list of 59 unique genes with an absolute correlation value above 0.5 contains 124 combinationsand contains many interesting correlations that connect cell shape with phenotype and potentially causal gene (available as supplementary data). To further corroborate this, we uploaded the Gene-Shape Correlation list into Cmap and retrieved the PCLs (Figure 7 a and b). Surprisingly no PCL signatures above absolute value 0.99 were associated with Area. In contrast, Minor Axis length has the largest number of associated signatures. Moreover, we have observed that PCLs *HIF activator, CDK inhibitor* and *Cell Cycle Inhibition GOF* were the most common signatures with the highest score. Interestingly, *ATPase inhibitor* was exclusively linked to shape parameter Euler Number, which was the most important for the prediction of the protein biosynthesis. At the same time, Euler Number has a very strong signature of the PCL *Protein Synthesis Inhibitor*. This example demonstrates that we can independently connect gene signalling and outcome in the phenotypic assay via cell shape features.

**Figure 7.**
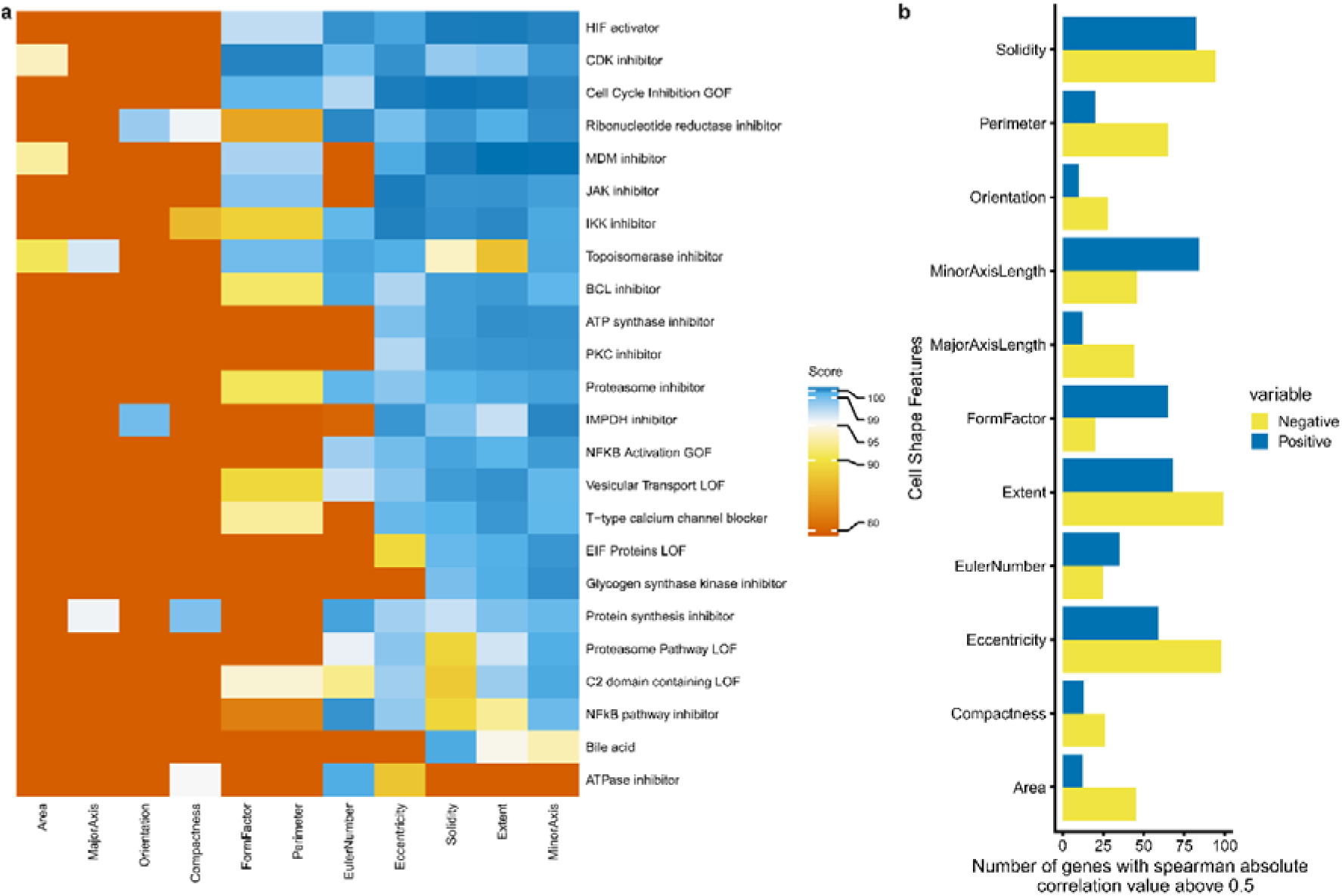
Cell biological processes related to cell shape features. **A.** Spearman correlation was calculated between gene expression and cell shape features. Genes with absolute Spearman correlation above 0.5 per condition (either per phenotype or per cell features) were used as input in the Connectivity Map, a gene expression database with more than 1 million profiles. All processes that have absolute score value above 99 at least for one condition are depicted, with 0 low and 100 as a high similarity. Biological processes have been ranked based on the number of conditions that can affect the process (specificity). **B.** Number of genes that were used for the analysis per shape feature.

### Comparison of different databases reveals a list of universal shape related genes

To investigate the broader relevance of our list of cell shape-correlated genes, we compared its overlap with two other data sets. In one study, whole transcriptome gene expression and related cell shape changes were induced by chemical compounds (Nassiri & McCall, 2018). In a second study, cell shapes were under the control of adhesive islands, and gene expression was assessed (Kilian, Bugarija, Lahn, & Mrksich, 2010). Overlap of all topographically-induced genes, 437 in total, and the two other gene sets yielded a list of only 12 genes (Figure 8a) and all the genes showed a strong Pearson correlation cell shape features (Figure 8b). Of the 12 genes found to be shape-predictable in all three studies, Minor Axis Length and Compactness correlated to eleven of them; Extent correlated to seven of them. As expected, Cell Orientation did not correlate to gene expression (Figure 8c). The above results demonstrate that filtering genes based on correlation to cell shape descriptors is a powerful method to find associations between gene expression, cell shape, and phenotype and that genes on the list of 275 genes can be considered as candidate genes directly influenced by cell shape. Indeed, of the twelve genes, seven have previously been directly linked to changes in cell shape:

**Figure 8:**
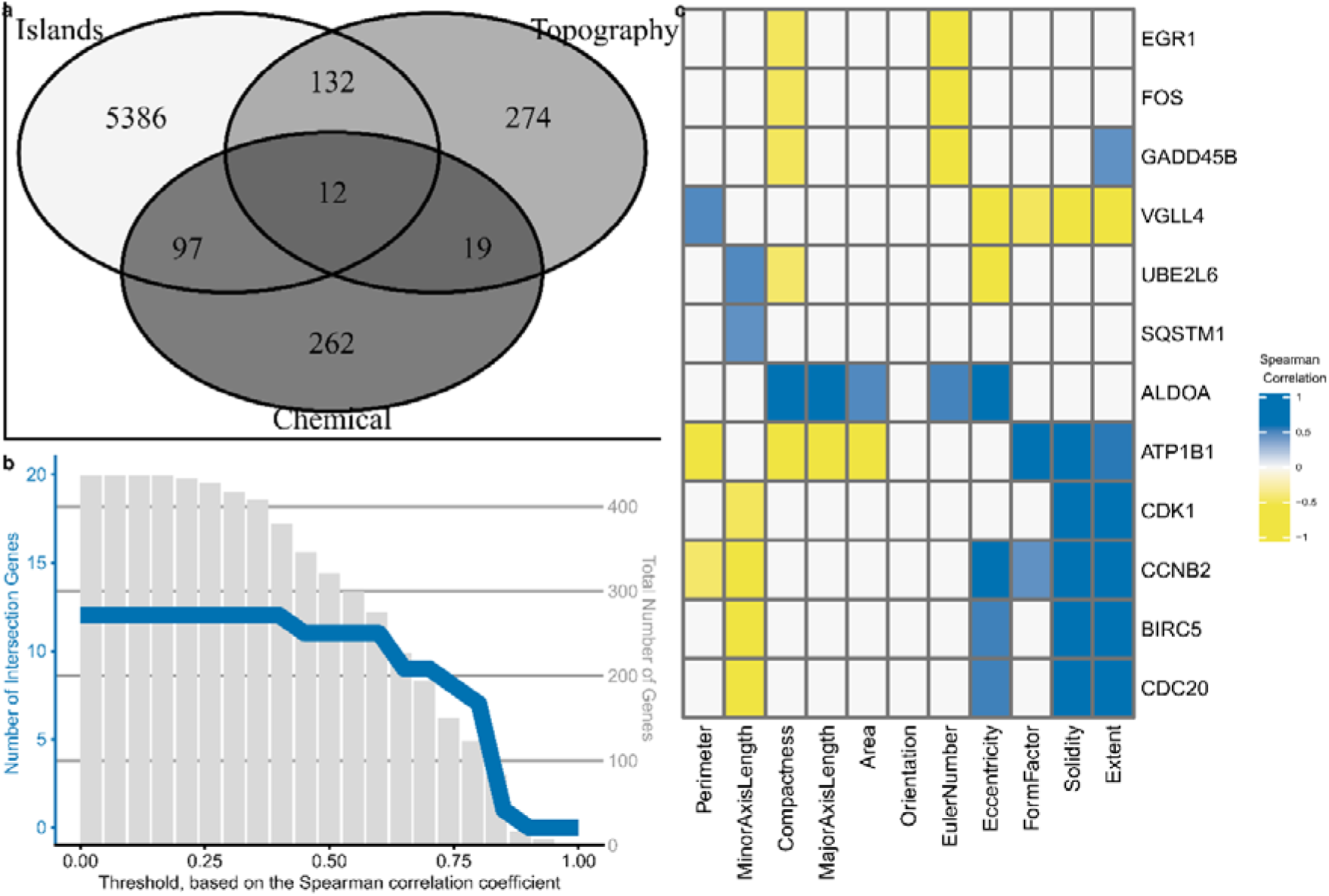
Genes related to shape are enriched in shape-based transcriptomics data sets. **A.** Venn diagram representing the overlap between genes differentially expressed on different adhesive islands, genes related to chemically induced shape changes and the 437 shape-based genes differentially expressed on the seven topographies with a fold change above 1.5. **B.** Filtering of the shape-specific genes based on the Spearman correlation score between the gene and at least on of the cell shape parameters. The red line and Y-axis at the left represents a number of selected shape-related genes with specified Spearman correlation threshold value (X-axis). Y-axis on the right represents the total number of genes that have Spearman correlation value above the specified threshold (X-axis). **C.** Heatmap that represents the correlation value between shape specific genes and shape parameters. All Spearman correlations with an absolute value below 0.4 are depicted for clarity.

- BIRC5 (Yap transcriptional target) (Muramatsu et al., 2010),
- EGR1 (Luxenburg, Amalia Pasolli, Williams, & Fuchs, 2011),
- FOS (Lamb et al., 1997),
- VGLL4 (YAP/TAZ inhibitor) (Moroishi, Hansen, & Guan, 2015),
- ALDOA (Du et al., 2014),
- SQSTM1 (cytoskeleton remodeling via autophagy) (Kadandale, Stender, Glass, & Kiger, 2010).

Three additional genes,

- CDK1 (Santamaría et al., 2007),
- GADD45B (Vairapandi, Balliet, Hoffman, & Liebermann, 2002) and
- CCNB2 (Fischer, Quaas, Steiner, & Engeland, 2015),

have been associated with proliferation. CDC 20 linked to both cell shape (Rho Signaling Protein) (David, Petit, & Bertoglio, 2012), and cells proliferation (Malumbres & Barbacid, 2009).

## Discussion and Conclusion

The molecular mechanisms connecting cell shape to basic cell functions and phenotype maintenance are important and yet remain largely unknown. Using high content imaging, transcriptomics and machine learning we were able to identify strong relationships between cell shape, molecular signalling and cellular phenotype. Some signal transduction pathways are particularly strongly correlated to topography-induced changes in cell shape. Changes in cell cycle-related signalling, and in the JAK, IKK and HIF pathways were observed in cells grown on most topographies. Other molecular signalling sigantures were more specific, such as SRC, ATPase inhibition and Glycogen synthesis kinase which were affected in MSCs grown on specific topographies. This confirms earlier findings by Nassiri and McCall who identified signatures of DNA damage and proliferation signalling changes in cell shape (Nassiri & McCall, 2018), and work of Jael et al. who noticed changes in NF-kB (Jain, Iyer, Kumar, & Shivashankar, 2013). Further support for the relevance of the genes and pathways discovered here in shape-related cell signalling comes from the overlap of 12 genes between our data set and those of two independent shape-related transcriptomics studies, and the known role of these genes. Thus, our systematic approach allowed us to confirm previous findings and to identify many novel unreported signatures. The data we have produced and publicly shared can be regarded as the first compendium of shape-gene expression and will likely lead to many new discoveries. The correlations can also be used to guide bio-instructive surfaces. For instance, more genes are affected by cell shape features such as Solidity and Extent then, for example, Compactness and some shape features are strongly correlated with certain signalling pathways. Thus, cell shape features can be used to manipulate cells’ molecular signatures with bio-active precision.

In this manuscript, we of obtained proof of principle for the correlation between image-based cell features and cell signalling and 11 basic cell shape features. There is likely an opportunity for mechanistic information within cell organelle shape and associated correlations with phenotype. Although a thorough investigation of this was beyond the scope of this work, we did discover a strong negative correlation between nuclear area and the expression of histone H3, suggesting a link between nuclear shape and epigenetic regulation (data not shown). The approach can be taken much further and feed into the creation of large-scale computational models which describe biomaterial surface design, cell shape and cell phenotype. This approach is referred to as Quantitative Structure-Activity Relationships (QSAR) and is widely used to model the biological activity of pharmaceutical compounds (Gramatica, 2007). It allows predicting the biological effect of untested compounds and thus narrowing down the search space.

To improve our shape-based models, we need more data which can be obtained by, e.g. upscaling the number of topographies from 28 in this manuscript to thousands, more extensive phenotypic characterisation of the cells for instance by using Cell Painting protocols (Bray et al. Nat Protocols) or other stains that display relevant biological processes and more extensive transcriptomics analysis on different cell types. As a start, data obtained in our project is freely available via our cBIT (Hebels, Carlier, Coonen, Theunissen, & de Boer, 2017) repository and can already be used to design a biomaterial that inhibits cell proliferation, for example, to reduce growth of the fibrosis tissue on the interface of the tissue-implant. The gene signatures related to cell shape and their underlying topographies can be used to browse the Connectivity Map database for genetic signatures that resemble the effect of small molecule or RNAi-induced gene signatures (Subramanian et al., 2017). This brings new opportunities to research the molecular pathways underlying shape-induced phenotypes. To accelerate the field of biomaterial engineering, this database should link information not only of topography design but also other material properties such as stiffness, ligand density, and any other relevant biomaterial properties. Based on systematic banking and mining of biomaterials, cell phenotype and transcriptomics data, we foresee a universal tool that will help understand and engineer the interface between cells and biomaterials. The work presented in this study outlines a method to search the large design space that links material properties to cell phenotype.

## Materials and Methods

### Cell culture

Human mesenchymal stem cells (hMSCs) were obtained from human bone marrow (donor d016), after informed consent as described previously (Mentik, 2013). Cells were cultured in basic medium (αMEM medium (Gibco, 22-571-038) supplemented with 10% fetal bovine serum (FBS) (Sigma-Aldrich batch number 013M3396) at 37 °C in a humid atmosphere with 5% CO^2^, unless stated differently. In all experiments, we used hMSCs at passage number four. Before cells seeding, surfaces were sterilized with 70% ethanol (Boom, 84010059.5000) and wetted in the basic medium during at least 2 hours.

### Adipogenesis

To induce adipogenesis, hMSCs (d016) were cultured for 3 weeks in adipogenic medium (DMEM (Gibco, 41-965-062), 100 U/ml penicillin plus 100 mg/ml streptomycin (Gibco, 15140-122), 10% fetal bovine serum (FBS) (Sigma-Aldrich batch number 013M3396), 0.2 mM indomethacin (Sigma-Aldrich, 57413), 0.5 mM IBMX (Sigma-Aldrich, I5879), 10^-6^ M dexamethasone (Sigma-Aldrich, D8893), 10 ug/ml insulin (human, Sigma-Aldrich, I9278). Cells were seeded in three replicas at density 15,000 cells/cm^2^ in 24 well plates, in basic medium. When the cells reached confluency on a flat surface, the culture medium was changed to adipogenic medium for 21 days, the medium was refreshed twice a week. To visualize lipids formation, cells were stained with oil red o (Sigma-Aldrich, O0625) as described before (De Boer, Wang, & Van Blitterswijk, 2004). Briefly, cells were fixed with 10 % formalin (Sigma-Aldrich, 501128) for 30 minutes at room temperature, rinsed with distilled water and washed with 60 % isopropanol (VWR, 20922.364). The sample was stained for 5 minutes in freshly filtered oil red o solution (stock: 500 mg oil red o (Sigma-Aldrich, O0625), 99 ml isopropanol, 1 ml distilled water; stain: 15 ml stock plus 10 ml distilled water).

### ALP induction

hMSCs (d016) were seeded on topographies at density 15,000 cells/cm^2^ in 96 well plates and cultured for 7 days in mineralization medium (basic medium supplemented with 2 mM L-glutamine (Gibco, 25030), 0.2 mM ascorbic acid (Sigma-Aldrich, A8960), 100 U/ml penicillin plus 100 mg/ml streptomycin (Gibco, 15140-122), 10^-8^ M dexamethasone (Sigma-Aldrich, D8893) plus 0.01 M β-glycerol phosphate (Sigma-Aldrich, 50020). Every surface had three replicas. The medium was changed every two days. Afterwards, cells were fixed with 4% Paraformaldehyde (PFA) (VWR, 30525), ALP was stained with immunofluorescent dye as described below.

### Cell Proliferation

hMSCSc (d016) were starved for 24 hours in αMEM (Gibco, 22-571-038) medium without FBS to synchronize their cell cycles. Then cells were trypsinized with trypsin-EDTA (0.05%) (Fisher Scientific, 25300054), trypsin was neutralized with the basic medium. Cells were seeded at a density of 10,000 cells/cm^2^ in triplicates and cultured in the presence of 1% FBS (Sigma-Aldrich batch number 013M3396). Cells were allowed to adhere overnight and afterwards, cells were cultured for 48 hours with 10 μM EdU component (Click-iT™ Plus EdU Alexa Fluor™ 594 Imaging Kit, Thermo Fischer Scientific) in the media. Next, incorporated EdU was imaged as described in the kit manual (Click-iT™ Plus EdU Alexa Fluor™ 594 Imaging Kit, Thermo Fischer Scientific).

### Protein Synthesis

To assess protein synthesis on the topographies we seeded hMSCs (d016) in triplicates at density 20,000 cells/cm^2^ on topographies in 96 well plates in basic medium and let them adhere for 24 hours. Later, the medium was replaced with DMEM high glucose, no L-glutamine, no L-methionine, no L-cystine, (Thermo Fischer Scientific, 21013024) supplemented with 580 mg/L L-glutamine (Thermo Fisher Scientific, 25030081), and 63 mg/L L-Cystine dihydrochloride (Sigma-Aldrich, C6727-25G). Prior cells fixation we added L-homopropargylglycine, L-methionine analogue (Thermo Fischer Scientific) for 1.5 hours. We further stained incorporated L-homopropargylglycine as described in the kit manual (Click-iT® HPG Alexa Fluor® Protein Synthesis Assay Kits, Thermo Fischer Scientific).

### Apoptosis

hMSCs (d016) were stained with dye for live cells, CellTracker™ Green CMFDA (Thermo Fischer Scientific, 11570166). During the next step, cells were seeded on topographies for 24 hours in a basic medium at density 10,000 cells/cm^2^ in 96 well plates. Next, we added a staurosporine (Abcam, ab120056), a protein kinase inhibitor, in the concentration of 0.5 uM and 250 ng/ml of annexin V (kind gift of Dr Leon J. Schurgers, Department of Biochemistry, University Maastricht) with a red fluorescent tag to detect apoptosis. To facilitate annexin V binding, the medium was supplemented with additional 0.7 mM of CaCl_2_ (VWR, 10043-52-4). Cells were monitored in an environmental chamber, where the temperature was maintained at 37 °C in a humid atmosphere with 5% CO_2_, during 66 hours, 16 images per well were captured every 25 minutes. Imaging was acquired on Nikon Ti Eclipse epifluorescent inverted microscope.

### Live imaging

hMSCs (d016) were stained with dye for live cells, CellTracker™ Green CMFDA (Thermo Fischer Scientific, 11570166). Afterwards, cells were seeded on topographies for 24 hours in a basic medium at density 10,000 cells/cm^2^ in 96 well plates. Cells were monitored in an environmental chamber, where the temperature was maintained at 37 °C in a humid atmosphere with 5% CO_2_, during 66 hours. 16 images per well were captured every 25 minutes. Imaging was acquired on Nikon Ti Eclipse epifluorescent inverted microscope.

### Staining and Imaging

Cells were fixed with 4% PFA (VWR, 30525) for ten minutes; next, samples were permeabilized with 0.01% Triton™ X-100 (VWR, 437002A) for 10 minutes with following blocking step in 0.5% bovine serum albumin (VWR, 421501J) at room temperature for 30 minutes. Samples were incubated with primary antibodies against ALP (RnD, mab1-148) overnight at 4 °C in 1:100 dilution. Labeling with secondary antibodies conjugated to fluorochrome Alexa Fluor® 647 goat anti-mouse IgG (Thermo Fisher Scientific, A21236) in dilution 1:300 and Alexa Fluor™ 568 Phalloidin (Thermo Fisher Scientific, A12380) in dilution 1:200 was made during 1 hour at room temperature, Next, after washing with PBS nuclear staining with 1:1,000 Hoechst 33342, trihydrochloride, trihydrate (Thermo Fischer, H1399) was done. Finally, the topographies were mounted on glass-bottom 24 well plates (Mobitec, 5231) or fluorocarbon-bottom 96 well plates (Mobitec, 5241) with ProLong® Diamond antifade mountant (Thermo Fischer Scientific, P36965) after 2 PBS (VWR, E404-200TABS) washes. Imaging was acquired on Nikon Ti Eclipse epifluorescent inverted microscope.

### Assesing cellular Morphology with the Cell Painting assay

hMSCs (d016) were seeded on topographies for 24 hours in basic medium at density 10,000 cells/cm^2^ in 24 well plates. Following the protocol of Bray et al., 30 minutes prior cells fixation 500 nM of MitoTracker™ deep red FM (Invitrogen, M22426) was added to cells. Afterwards, cells were washed with HBSS (Fisher Scientific, 11540476) and fixed with 4% PFA (VWR, 30525) for 20 minutes at room temperature. Samples were permeabilized with 0.1% Triton™ 100 (VWR, 437002A) for 10 minutes with the following washing with HBSS (Fisher Scientific, 11540476). We further added wheat germ agglutinin Alexa Fluor™ 594 conjugate (Invitrogen, W11262) at concentration 5 µg/ml, concanavalin A, Alexa Fluor™ 488 conjugate (Invitrogen, C11252) at concentration 50 µg/ml, SYTO™ 14 green fluorescent nucleic acid stain (Invitrogen, S7576) at concentration 10 µM, 1:40 Alexa Fluor™ 568 Phalloidin (Thermo Fischer Scientific, A12380) and 1:2,000 Hoechst 33342, trihydrochloride, trihydrate (Thermo Fischer, H1399) for 30 minutes. Finally, the topographies were mounted on glass-bottom 24 well plates (Mobitec) with ProLong® Diamond Antifade Mountant (Thermo Fischer Scientific, P36965) after 2 PBS washes.

### Transcriptional Profiling

hMSCs (d016) were seeded on selected topographies for 24 hours in a basic medium at density 15,000 cells/cm^2^ in 24 well plates in three replicas, total RNA was isolated using the Nucleospin RNA isolation kit (Macherey–Nagel). Then, from 100 ng of RNA, cRNA was synthesized using the Illumina TotalPrep RNA amplification kit, and both RNA and cRNA quality was verified on a Bioanalyzer 2100 (Agilent). Microarrays were performed using Illumina HT-12 v4 expression Beadchips. Briefly, 750 ng of cRNA was hybridized on the array overnight, after which the array was washed and blocked. Then, by addition of streptavidin Cy-3, a fluorescent signal was developed. Arrays were scanned on an Illumina Beadarray reader and raw intensity values were background corrected in BeadStudio (Illumina). Further data processing and statistical testing were performed using the online portal arrayanalysis.org (Eijseen 2015). The probe-level raw intensity values were quantile normalized and transformed using variance stabilization (VSN). A detection threshold of 0.01 was used to reduce the number of false positives. A linear modelling approach with empirical Bayesian methods, as implemented in Limma package (Smith, 2005), was applied for differential expression analysis of the resulting probe-level expression values. P-values were corrected for multiple testing using the Benjamini and Hochberg method (Benjamin and Hochberg 2015). Genes were considered differentially expressed when a corrected p-value below 0.05 was reached. Pathway over-representation analysis was performed using the web tool ConsensusPathDB (CPDB), which provides a comprehensive pathway analysis covering most public resources for interactions (Kamburov et al., 2010). Over-representation analysis was performed on a set of differentially expressed genes and a background list containing all measured genes was used to improve the statistical evaluation of the pathways. Pathways with a false discovery rate-corrected p-value <0.05 were considered significant.

Network analysis was carried out in two steps. CPDB contains an induced network module which uses the interactions described in all the public resources to build a network based on a list of input genes. At first, a network was generated on the same list of differentially expressed genes as used for pathway analysis using a z-score threshold of 20. Only binary protein interactions of low, medium and high confidence were selected and intermediate genes were allowed to be added to the network in order to improve inter-gene connectivity. The resulting network was subsequently imported into CytoScape and the plugin CyTargetLinker was used to extend the CPDB network by adding transcription factors from the transcription factor target database TFe (Transcription Factor encyclopedia). TFe is a smaller scale manual literature curation project containing 1531 human transcription factor target interactions respectively.

### Selection of Surfaces and Clustering

To be able to find cell shapes that were repeatedly induced by topography and remove any imaging artefacts or biological variation we performed the following filtering steps. Among other outlier detection tools we used, so called, 1.5 interquartile range (IQR) rule. It works as following: calculate 1^st^ and 3^rd^ quantile of the data (value of the first 25% and 75% of ordered data, correspondingly), difference between 3^rd^ and 1^st^ quantiles is IQR, all the data points that lie outside of the following range: 1^st^ quantile-1.5*IQR, 3^rd^ quantile + 1.5*IQR, considered being outlines. First, we removed replicas based on cell density. We performed this on a per-surface basis, to avoid removal of the surfaces with a unique response. Replicas with too low or to high cell number were discarded using the 1.5 IQR rule. Afterwards, we removed cells that were outliers in terms of area and perimeter. Outliers were removed sequentially, first based on area and then based on perimeter values using the 1.5 IQR rule. We further employed Moutlier package in R (Gupta, Gao, Aggarwal, & Han, 2014) to remove outliers based on cell shapes. This approach allows finding outliers by using information from multiple features simultaneously. We did this by looking at the distribution of cell shapes in a 5-dimensional shape space and then removing outliers using the Mahalanobis-based outlier detection. These 5 shape features were selected from 11 features that have correlation index less than 0.7 between each other. Finally, for each surface, we retained only those replicates that were of good quality, those that were reproducible. The reproducibility was measured using the correlation of cell shapes between replicas. We excluded replicas that have correlation index less than 0.5, in comparison to all the replicas. Next, we summarized cell shape features per surface by taking median features of all the cells. Afterwards, we removed redundant surfaces, those that were highly correlated (Pearson correlation index above 0.95) to each other based on cell shape features. To reduce the dimensionality of the data further, we performed principal components analysis. We have chosen first seven principal components that captured the most variations of the data. We further used hierarchical clustering analysis with Ward linkage to find clusters of cell shapes. A distance matrix was calculated with Euclidean distances.

### Image and Data Analysis

Open-source software Cell Profiler 2.1 (CP) was used for image analysis (Carpenter et al., 2006). In order to perform automated image analysis in CP, a robust pipeline able to recognize different cell features was built.

The t-SNE analysis was performed with perplexity equals to 6 in package Rtsne (Krijthe, van der Maaten, & Krijthe, 2018). The principal component analysis was performed with “prcomp” function on scaled and centred data.

### Training the Model

We employed Lasso regression model from Glmnet package in R (Hastie & Qian, 2014) to find cell shape features that can predict the response of the phenotypic assays. The model was trained with Leave One Out Cross validation approach. We assessed the performance of the model by reporting the Pearson correlation index (r) between predicted and observed values on the training data. To show the goodness of fit we also reported R^2^. The hyperparameter lambda was optimized separately for every model. Cells on flat control surfaces were excluded because of their distinct response, in comparison to the rest of topographies, which would bias the model.

### Connectivity Map Analysis

List of genes was uploaded into Connectivity Map (Subramanian, Narayan, et al. 2017), a library of gene expression signatures induced by chemical compounds or genetic interference and a connectivity score between the topography-induced DEGs and Connectivity Map DEGs were retrieved. We focused on perturbant classes (PCL), which are groups of signatures in which members belong to the same gene family or are targeted by the same compound. To find cell shape signatures we have uploaded genes that have absolute correlation value above 0.5. Where genes with positive correlation values were loaded as “upregulated genes” and genes with negative correlation values were loaded as “downregulated genes. The results were futher exported to csv file and vizualized in R.

### TopoChip and enlarged surfaces fabrication

The complete design of the TopoChip is described in detail elsewhere (Unadkat et al., 2011). In summary: The in silico design of the TopoChip is generated using an algorithm that combines the primitive shapes: triangle, circle and rectangle to generate complex shapes. The primitive shapes “triangle, circle, rectangle” were chosen because the stochastic combination of these shapes in varying orientation, numbers and sizes result in the generation of nearly any possible shape. The *in silico* design was used to create a chromium mask for photolithography.

Both the TopoChips and enlarged surface areas of hit topographies were prepared by hot embossing of (poly)styrene (PS) films (Goodfellow).

Briefly, standard photolithography and deep reactive etching were used to produce the inverse structure of the topographies on a silicon wafer. The silicon master mould was then used to make a positive mould in poly(dimethylsiloxane) (PDMS). Subsequently, a second negative mould in OrmoStamp hybrid polymer (micro resist technology Gmbh) was obtained from the PDMS mould. This mould serves as the template for hot embossing the TopoChips and enlarged topographically enhanced surface areas in 190 µm thick PS films using 140 °C and 10 barr for 5 minutes. Prior to cell culture, the topographically enhanced PS films were O_2_-plasma treated to increase protein attachment to the substrate surface in order to increase cell attachment.

### Statistical analyses and Data Visualization

Statistical analysis was performed in R ver. 3.2.5 (Ihaka & Gentleman, 1996), graphs were generated in R package ggplot2 (Wickham, 2016) and cowplot (Wilke, 2016).

## Acknowledgements

We thank Prof. Dr. Kris Kilian for providing microarray data. The research leading to these results has received funding from the European Union’s Seventh Framework Programme (FP7/2007-2013) under grant agreement no 289720. AV, SV, MK, NB, AC and JdB acknowledge the financial contribution of the Province of Limburg. AC gratefully acknowledges her VENI grant (number 15057) from the Dutch Science Foundation (NWO).

