## Supplementary figures and images for "On the correlation between material-induced cell shape and phenotypical response of human mesenchymal stem cells"

### Supplementary Figure 1

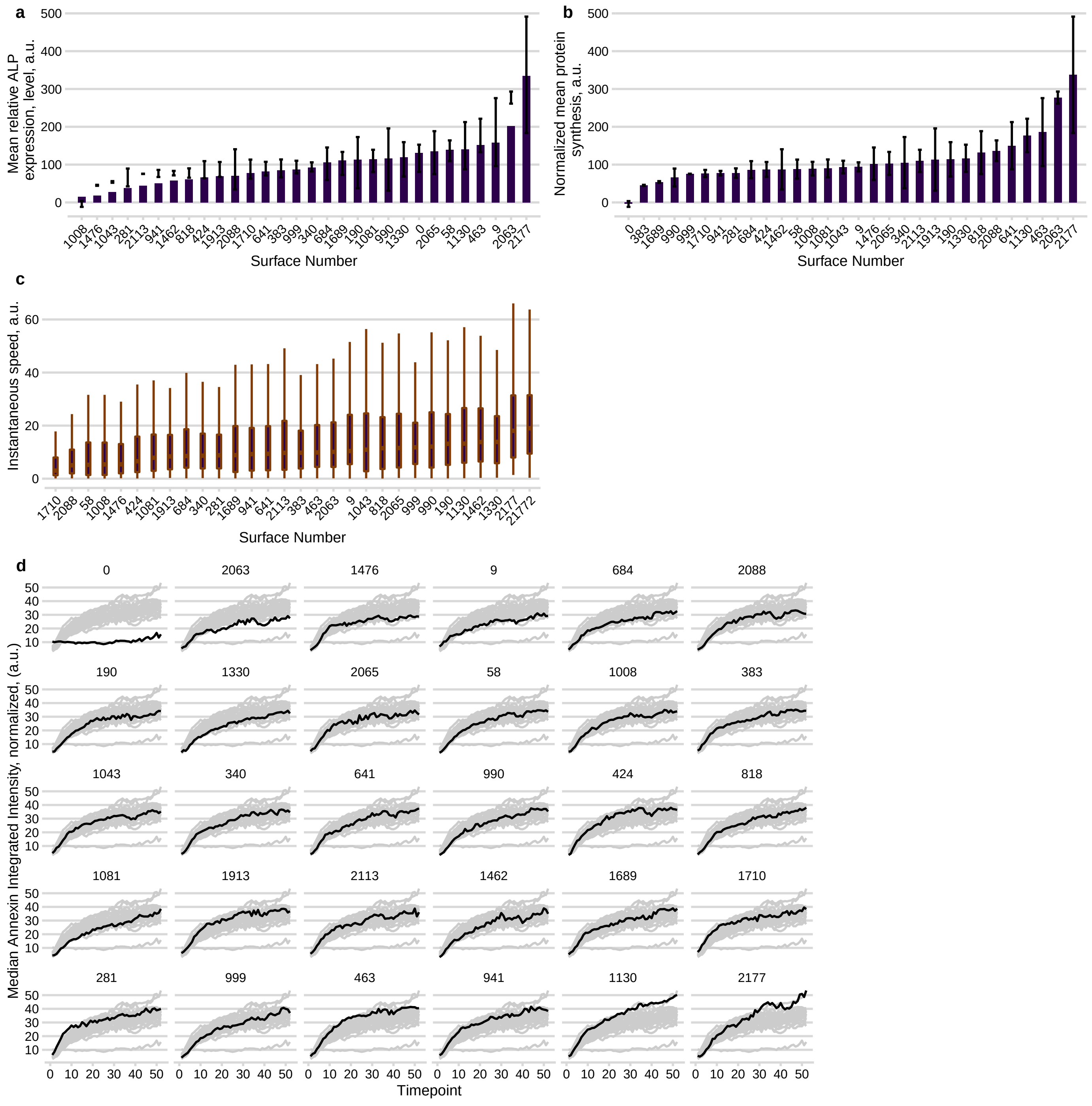

### Supplementary Figure 2

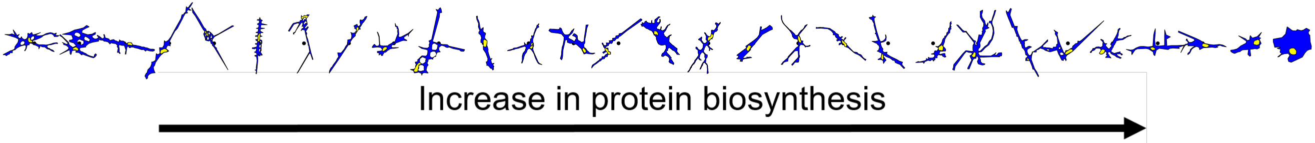
