## Supplementary material for "On the correlation between material-induced cell shape and phenotypical response of human mesenchymal stem cells": tables

**Table 1. List of basic cells shape features and their definitions.**

| **Cell Shape Feature** | **Definition** |
| --- | --- |
| Area | The actual number of pixels in the region |
| EulerNumber | The number of objects in the region minus the number of holes in those objects, assuming 8-connectivity. |
| MajorAxisLength | The length (in pixels) of the major axis of the ellipse that has the same normalized second central moments as the region. |
| MinorAxisLenght | The length (in pixels) of the minor axis of the ellipse that has the same normalized second central moments as the region. |
| Orientation | The angle (in degrees ranging from -90 to 90 degrees) between the x-axis and the major axis of the ellipse that has the same second-moments as the region |
| Perimeter | The total number of pixels around the boundary of each region in the image |

**Table 2. List of elaborate cells shape features, their definitions and schematic.**

| **Feature** | **Definition** | **Schematic** |
| --- | --- | --- |
| Compactness | The variance of the radial distance of the object's pixels from the centroid divided by the area. | 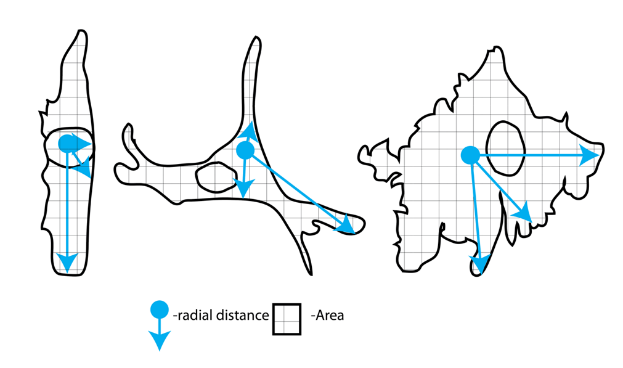 |
| Eccentricity | The eccentricity of the ellipse that has the same second-moments as the region. The eccentricity is the ratio of the distance between the foci of the ellipse and its major axis length. The value is between 0 and 1. (0 and 1 are degenerate cases; an ellipse whose eccentricity is 0 is actually a circle, while an ellipse whose eccentricity is 1 is a line segment.) | 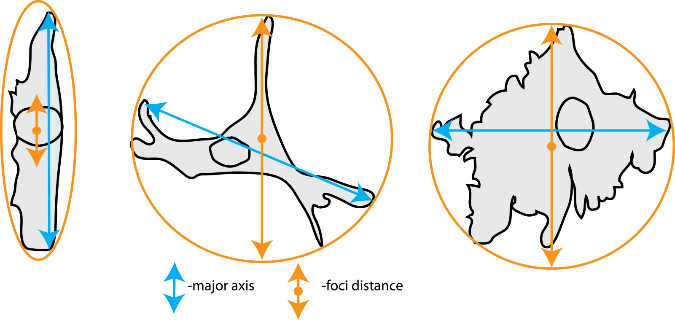 |
| Extent | The proportion of the pixels in the bounding box that are also in the region. Computed as the Area divided by the area of the bounding box. | 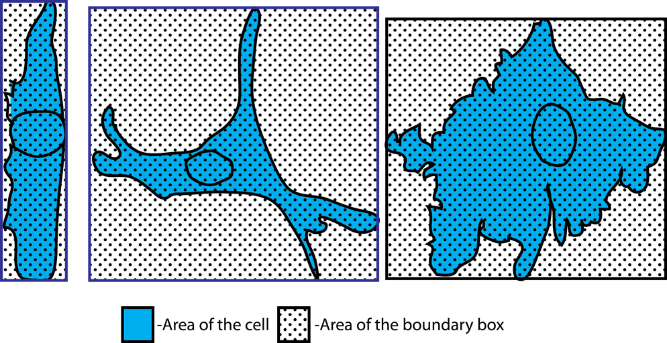 |
| Form Factor | Calculated as 4*π*Area/Perimeter^2^. Equals 1 for a perfectly circular object | 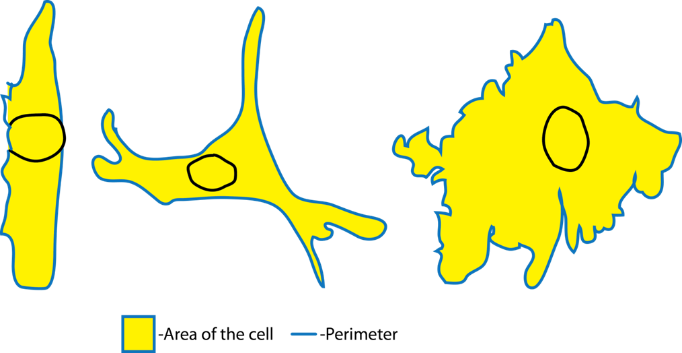 |
| Solidity | The proportion of the pixels in the convex hull that are also in the object, i.e. ObjectArea/ConvexHullArea. Equals 1 for a solid object (i.e., one with no holes or has a concave boundary), or <1 for an object with holes or possessing a convex/irregular boundary. | 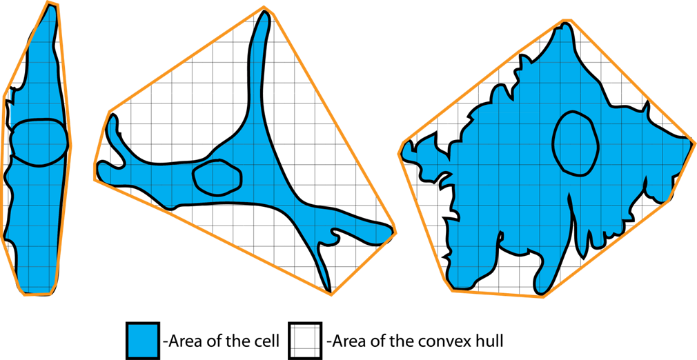 |
